# The structure of the UDP-Glc/GlcNAc 4-epimerase from the human pathogen *Campylobacter jejuni*

**DOI:** 10.1101/2020.09.22.308395

**Authors:** Hyun Gi Yun, Kyoung-Soon Jang, Shiho Tanaka, William M. Clemons

## Abstract

Worldwide, the food-born pathogen *Campylobacter jejuni* is the leading bacterial source of human gastroenteritis. *C. jejuni* produces a variety of diverse cell-surface carbohydrates that are essential for pathogenicity. A critical component of these oligo- and polysaccharides is the sugar *N*-acetylgalactosamine (GalNAc). The sole source of this sugar is the epimerization of UDP-*N*-acetylglucosamine (GlcNAc), a reaction catalyzed by the enzyme UDP-GlcNAc 4-epimerase (Gne). This enzyme is unique among known bacterial epimerases in that it also catalyzes the equivalent reaction with the non-*N*-acetylated sugars. Understanding how *Cj*Gne catalyzes these various interconversions is critical to designing novel inhibitors of this enzyme. Here, to further the mechanistic understanding we present a 2.0Å structure of *Cj*Gne with its NAD^+^ co-factor bound. Based on novel features found in the structure we perform a variety of biochemical studies to probe the mechanism and compare these results to another bifunctional epimerase, human GalE. We further show that ebselen, previously identified for inhibition of *Hs*GalE, is active against *Cj*Gne, suggesting a route for antibiotic development.

## Introduction

*Campylobacter jejuni*, a microaerophilic pathogen, is a commensal of chickens and other avians and the most prevalent member of the *Campylobacter spp*., which are the leading causes of bacterial gastroenteritis worldwide (1, 2). It can be associated with post-infectious sequelae include Guillain-Barrè syndrome (3), bacteremia (4), and reactive arthritis (5). The *C. jejuni* glycome contains a number of surface-accessible carbohydrate structures required for interactions with the various hosts that include capsular polysaccharide (CPS), lipooligosaccharide (LOS), and N- and O-linked glycans (6–12). Variability in the nature of the surface glycans challenges the development of anti-*Campylobacter* therapies (13, 14).

All of the exposed currently identified glycans in *C. jejuni* contain a GalNAc residue (14–17), which is only produced through the epimerization of GlcNAc by the enzyme encoded by the gene *gne* (UDP-<u>G</u>lc<u>N</u>Ac 4-<u>e</u>pimerase). It was recently reannotated from *galE* (UDP-<u>Gal</u> 4-<u>e</u>pimerase) due to the discovery of the UDP-GlcNAc epimerization activity (16) (Figure 1A). In the *C. jejuni* genome (strain NCTC11168), this gene is located between the gene clusters of the N-linked glycosylation and LOS biosynthesis pathways (18). The gene in *C. jejuni* was first believed to be involved in the LPS biosynthesis pathway (10), but later its functional role in N-linked glycosylation was elucidated (19, 20). Furthermore, the experiments with an insertional mutant of *gne* indicated that it is responsible for providing GalNAc residues to the three major cell-surface glycoconjugates (16). The bifunctional protein *Cj*Gne represents a potential therapeutic target as most oligosaccharides in *C. jejuni*, which contain a GalNAc residue, are required for pathogenesis (21). A mechanistic understanding is key to developing therapeutics that target this unusual epimerase.

**Figure 1.**
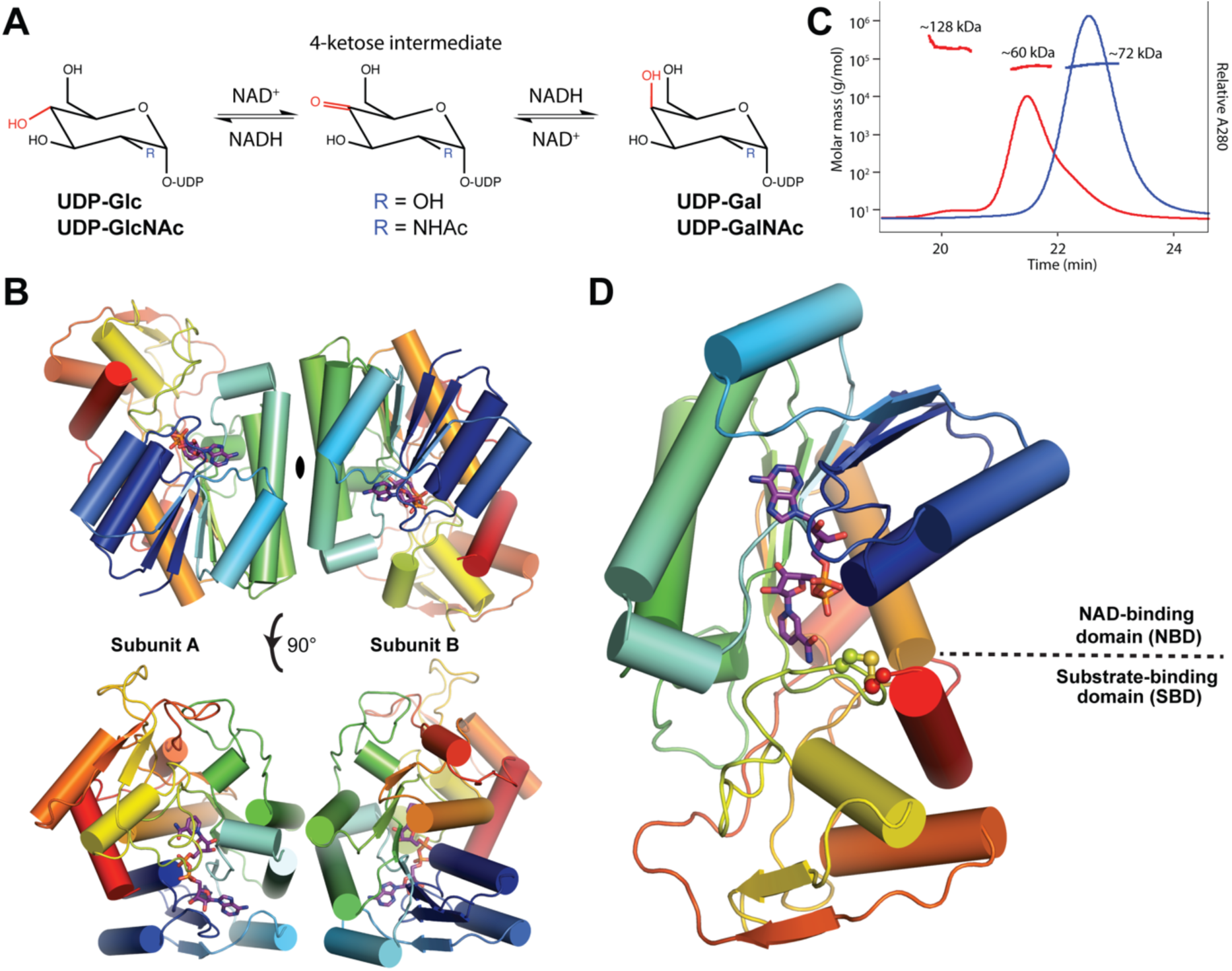
The 2.0Å crystal structure of *Cj*Gne in complex with NAD^+^. **A**. A catalytic reaction scheme of *Cj*Gne involving four substrates, UDP-Glc, UDP-Gal, UDP-GlcNAc, UDP-GalNAc, and 4-ketose intermediate. **B**. (Top) An asymmetric unit contains two molecules of *Cj*Gne with a two-fold symmetry. (Bottom) The two molecules are rotated by 90°. In both cases, a monomer in the cartoon representation is colored with rainbow; N-terminus is with red and C-terminus is with blue. One NAD^+^ molecule in each monomer is shown in stick representation with the purple backbone. **C**. The SEC-MALLS result shows *Cj*Gne forms a dimer (∼72 kDa; blue) in solution as compared to the bovine serum albumin (BSA) standard (monomer: ∼60 kDa; dimer: ∼128 kDa; red). **D**. A monomer is colored the same as in Figure 1B. Additionally, the disulfide bond is depicted in yellow spheres. NBD and SBD are also labeled.

UDP-hexose 4-epimerases, including *Cj*Gne, belong to a protein family of short-chain dehydrogenases/reductases (SDRs) (22). SDR enzymes possess diverse substrate specificity (23), as seen in the UDP-hexose 4-epimerases that have undergone detailed characterization (24). Structures of UDP-hexose 4-epimerases are available from all three domains of life: GalE from *Escherichia coli* (*Ec*GalE) (25–27), GalE from *Trypanosoma brucei* (*Tb*GalE) (28), GalE from *Pyrobaculum calidifontis* (*Pc*GalE) (29), GalE from *Homo sapiens* (*Hs*GalE) (30, 31), GalE from *Thermotoga maritima* (*Tm*GalE) (32), WbpP from *Pseudomonas aeruginosa* (*Pa*WbpP) (33), and WbgU from *Plesiomonas shigelloids* (*Ps*WbgU) (34). A classification scheme has been proposed for substrate preference of the UDP-hexose 4-epimerases (33). The scheme categorizes the epimerases in three different groups depending on the side chain size of six key active site residues. Group 1 epimerases, such as *Ec*GalE and *Tb*GalE, interconvert only the non-acetylated moieties, UDP-Glc and UDP-Gal. Group 3 enzymes prefer epimerizing acetylated moieties and the examples are *Pa*WbpP and *Ps*WbgU. Group 2 members include *Cj*Gne and *Hs*GalE with no preference for interconversion of the non-acetylated or acetylated UDP-hexoses. All of the UDP-hexose 4-epimerases structurally characterized so far are functional either in the Leloir pathway for galactose metabolism or the LPS O-antigen biosynthesis pathway. No epimerase known to act on multiple pathways has yet been a subject of structural studies.

Toward a mechanistic understanding of this bi-functional multi-pathway enzyme, here we provide a 2.0Å crystal structure of NAD^+^-bound *Cj*Gne. Structural characteristics of *Cj*Gne that are common or distinct to its homologs are discussed. Based on the structural features and the epimerization results of the wild type and its mutants, we propose some critical residues of *Cj*Gne that are catalytically or structurally important for shaping its substrate-binding site and determining substrate specificity.

## Results

### The overall architecture of *Cj*Gne bound with NAD^+^

The gene for *Cj*Gne was cloned into an expression vector, induced in *E. coli*, purified by chromatography, and crystallized via conditions initially obtained through standard screens. Crystals diffracted to 2.0Å and a complete data set was collected. Data were processed using standard tools and the structure was solved by molecular replacement using *Ba*GalE (PDB entry: 2C20). The fully refined model contained a dimer of *Cj*Gne with an R-factor of 19.5% and an R_free_ of 22.5%. A complete atomic model containing residues 2-328 was attained for each with an RMSD of 0.38 Å between the two copies. Crystallographic statistics are found in Table 1. The two copies in the asymmetric unit are related by a two-fold rotational axis (Figure 1B). Consistent with this as a dimer interface, in solution the enzyme purified as a dimer which was verified using SEC-MALLS. The apparent molecular weight of 72 kDa was comparable to a dimer of 74 kDa (Figure 1C). For clarity, subunit A (indicated in Figure 1B) will be used in figures when only one is present.

**Table 1.** Statistics of x-ray data collection and refinement.

| **Data collection** |  |
| --- | --- |
| Space group | P4<sub>1</sub> 2<sub>1</sub> 2 |
| Cell dimensions |  |
| a/b/c (Å) | 87.7/ 87.7/ 261.66 |
| $\alpha/\beta/\gamma$ (°) | 90/ 90/ 90 |
| Resolution (Å)<sup>a</sup> | 65.42–2.04 (2.09–2.04) |
| R<sub>merge</sub><sup>a</sup> | 0.052 (0.677) |
| I/$\sigma$ (I)<sup>a</sup> | 18.0 (2.4) |
| Completeness (%)<sup>a</sup> | 99.9 (100) |
| Redundancy<sup>a</sup> | 6.6 (6.8) |
| **Refinement** |  |
| Resolution (Å) | 62.0 – 2.0 |
| No. reflections | 65,906 |
| R<sub>work</sub>/ R<sub>free</sub> | 19.5/ 22.5 |
| No. atoms | 5,463 |
| Protein | 5,290 |
| Ligand/ion | 6 |
| Water | 167 |
| Mean B-factor | 38.1 |
| RMS<sup>b</sup> deviations |  |
| Bond lengths (Å) | 0.008 |
| Bond angles (°) | 1.1 |

<sup>a</sup> Values in parentheses are for highest-resolution shell<sup>b</sup> RMS, root mean square

The general structure of *Cj*Gne is consistent with that seen for related epimerases with a Glycosyl-transferase A fold. Each subunit of *Cj*Gne contains two domains and one NAD^+^ cofactor (Figure 1D). The domain that contains the N-terminus (M1-Y174, I230-H260, I295-D312) is composed of a central, twisted, parallel, seven–stranded, β–sheet flanked on each side by four α-helices. This domain resembles a ‘Rossmann-fold’ motif where the NAD^+^ binds that is common to SDR enzymes (35) and will be referred to as the NAD-binding domain (NBD). The substrate-binding domain (SBD) containing mostly C-terminal residues (F175-F229, G261-L294, D313-C328) is composed of three α-helices and two parallel β-strands. Previous structural studies of related epimerases demonstrate that the UDP-hexose substrate resides in the cleft between the two domains with the UDP moiety primarily contacting the SBD (26).

### The conserved NAD-binding domain (NBD)

Both NAD^+^ and NADH have been previously resolved in crystal structures of related epimerases (27). The only structural difference between NAD^+^ and NADH is in the nicotinamide ring (Figure 2A). At this resolution, analyzing the 2Fo-Fc map around the nicotinamide ring did not conclusively demonstrate whether or not the ring is planar (Figure 2B). We instead compared Fo-Fc maps after refining the data with the model of NAD^+^ and NADH. When NADH was used in refinement an additional positive density in the Fo-Fc map appeared in the nicotinamide ring; no density was seen for NAD^+^ (Figure 2B). The unaccounted-for electron density with NADH is consistent with the additional electrons found in the nicotinamide ring of NAD^+^, which we conclude is the major state in our crystal form. The nicotinamide ring adopts the *syn* conformation with respect to the ribose as seen in some of the *Ec*GalE (PDB ID: 1NAI, 1XEL, 1LRL) and all of the *Hs*GalE structures (PDB ID: 1EK5, 1EK6, 1HZJ) published. However, it is unclear whether there is any correlation between the identity and conformation of this cofactor in the NAD-binding domain and substrate identity.

**Figure 2.**
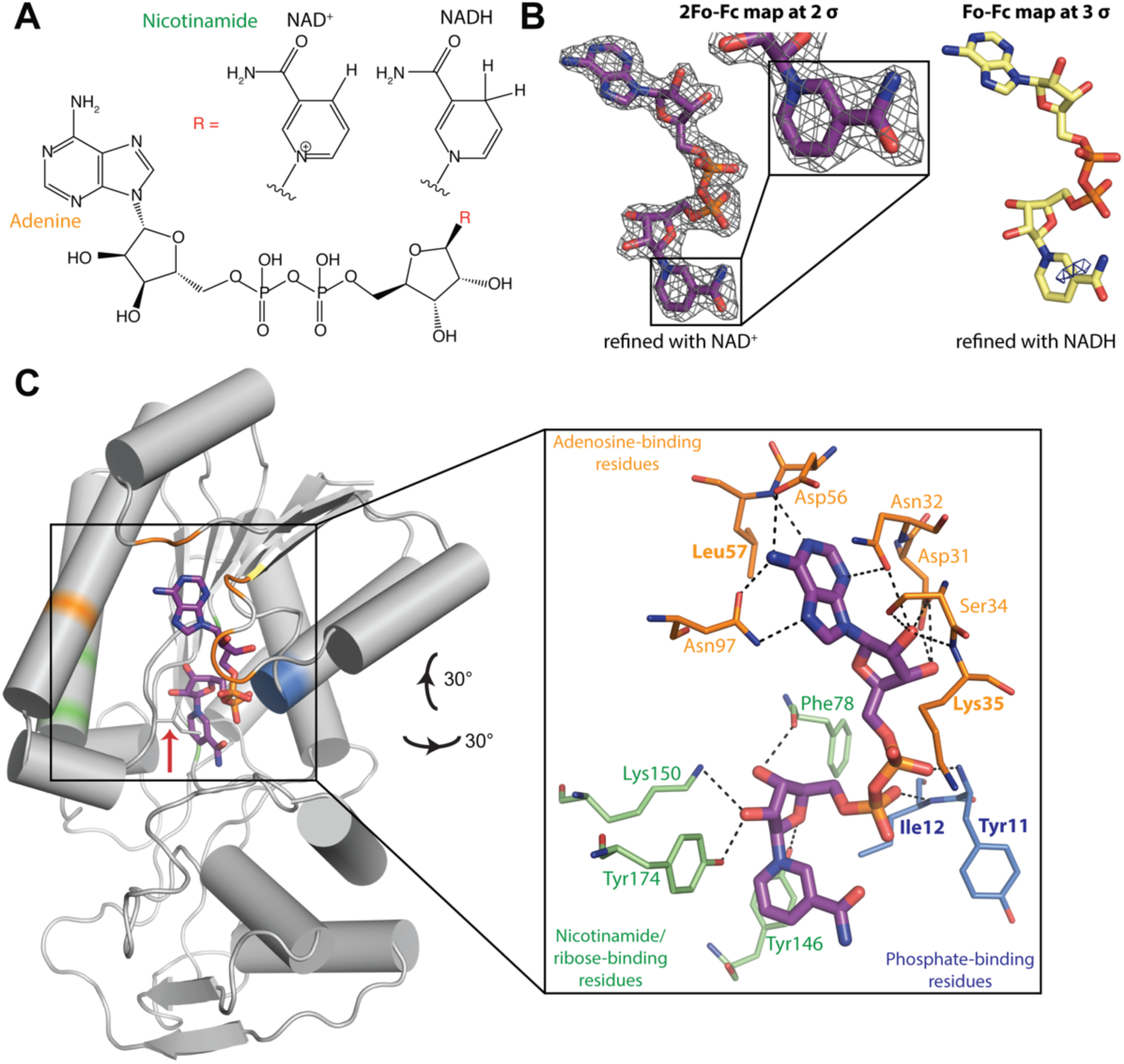
NAD^+^ and NBD. **A**. Comparison of the chemical structures of NAD^+^ and NADH. The only difference lies in the nicotinamide ring, depicted as a R group. **B**. On the left, the 2Fo-Fc map that is refined with NAD^+^ is shown in mesh at 2 σ and the magnified view of the nicotinamide ring. On the right, the Fo-Fc map that is refined with NADH is shown in mesh at 3 σ. **C**. On the left, a monomer of *Cj*Gne in the same orientation as in Figure 1D is colored grey. NAD^+^ is shown in stick representation and its backbone is purple. Ile82 is indicated by a red arrow. On the right, this is a magnified view of the NBD. The residues that are in a hydrogen-bonding distance to adenosine, phosphates, and nicotinamide/ribose are shown in stick representation and colored orange, blue, and green, respectively. The residues that interact with NAD^+^ via backbone are bolded (Tyr11, Ile12, Lys35, and Leu57). Hydrogen bonds are shown in black dashed lines.

The interaction with NAD^+^ seen here is characteristic of related NAD-binding domains. The details were first described in the structure of GalE from *E. coli* where a series of conserved residues line the binding pocket (PDB ID: 1NAI) (36). In *Cj*Gne, Ser34 and Lys35 forms one more and one less hydrogen bond than the corresponding residues in *Ec*GalE, respectively (Figure 2C). In *Ec*GalE, an additional residue Lys84 stabilizes the pyrophosphate group of NAD^+^ via two hydrogen bonds, while the corresponding residue Ile82 in *Cj*Gne makes none (Figure 2C). Overall, seven residues (Asp31, Asn21, Ser34, Lys35, Asp56, Leu57, Asn97) are in hydrogen-bonding distance with the adenosine of NAD^+^. Two residues (Tyr11, Ile12) and four residues (Phe78, Tyr146, Lys150, Tyr174) are interacting through hydrogen bonds with the pyrophosphate group and nicotinamide/ribose of NAD^+^, respectively.

### Apo-substrate-binding pocket is expanded

In a comparison to the other bifunctional epimerase *Hs*GalE, our apo-*Cj*Gne structure aligns best with the available apo-GalE ‘resting enzyme’ from human (PDB ID: 1EK5) (30) with an RMSD of 1.292 Å. The SBDs alone align with the RMSD value is 1.224 Å, similar to the overall RMSD. In order to find differences in the substrate-binding sites, *Cj*Gne was aligned only with the SBD of *Hs*GalE bound to UDP-GlcNAc (PDB ID: 1HZJ) (Figure 3A). Based on the superposition location of the substrate, while R300 and D303 of *Hs*GalE stabilize UDP-GlcNAc through some hydrogen bonds (Figure 3B), the corresponding residues (R287 and D290) of *Cj*Gne are unlikely to contact the UDP-GlcNAc (Figure 3C). Instead, D290 forms a hydrogen bond with R287, which in turn interacts with the backbone of Y190 and P191 (Figure 3C and 3D). Interestingly, Y190 and P191 are part of the shifted loop in *Cj*Gne that will be described in the next section.

**Figure 3.**
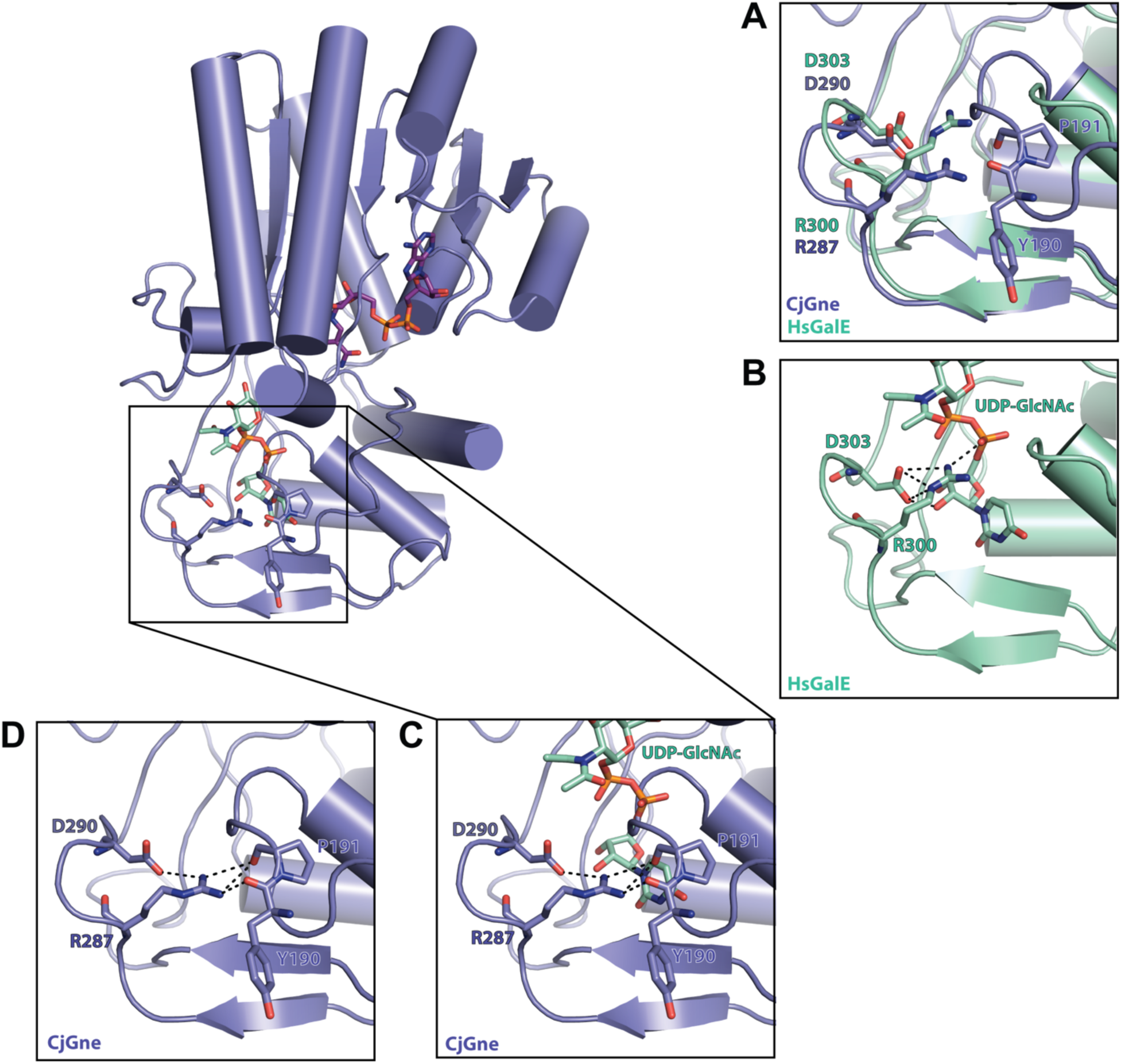
SBD of *Cj*Gne. A *Cj*Gne monomer is shown in complex with NAD^+^ and UDP-GlcNAc from *Hs*GalE (PDB ID: 1HZJ). **A**. Aligned *Cj*Gne (slate) and the *Hs*GalE SBD (green cyan) in the box region. Y190, P191, R287, and D290 of *Cj*Gne and R300 and D303 of *Hs*GalE are shown in stick representation. **B**. Same region as in Figure 3A, but only *Hs*GalE is shown along with the polar contacts in black dashed lines. **C**. Same region as in Figure 3A, but *Cj*Gne and UDP-GlcNAc from *Hs*GalE are shown with polar contacts in black dashed lines. **D**. Same region as in Figure 3A, but only *Cj*Gne is shown with polar contacts between R287/D290 and Y190/P191.

### A shifted loop in the *Cj*Gne structure

Our crystal structure of *Cj*Gne revealed a surprising feature not found in any other structures of its related enzymes. In *Cj*Gne, one loop region (174-YF-NAGACMDYTLGQRYPKATL-195) is shifted toward the NBD obstructing the substrate-binding pocket (Figure 4A). In the crystal, this shift is locked by a disulfide bond between C181 and the C-terminal residue, C328 (Figure 4A). Both of these cysteine residues are not found in related epimerases, except for closely related *Campylobacter spp* where they always occur as a pair (Figure 4B, 4C, and 4D). We presume disulfide bond formation results in stabilization of the shifted loop. *Cj*Gne localizes in the cytoplasm and, under typical media conditions, this would be a reducing environment incompatible with a disulfide bond. During purification fresh DTT was added to all buffers (pH 7.5) to maintain a reducing environment. However, the presence of the disulfide suggested this was insufficient. DTT is reported to be readily oxidized above pH 7.5 (37) and in the presence of metal ions like Ni^2+^ contaminants from Ni column (38). TCEP was added as an alternative to the final protein solution and incubated overnight before setting up crystal trays. It was also added to cryo-protecting conditions when harvesting crystals. Although TCEP is known to be stable at both acidic and basic conditions (37) and active with some metal contaminants, all acquired crystals had the disulfide bond. This suggests that the formation of the disulfide is favored by the protein.

**Figure 4.**
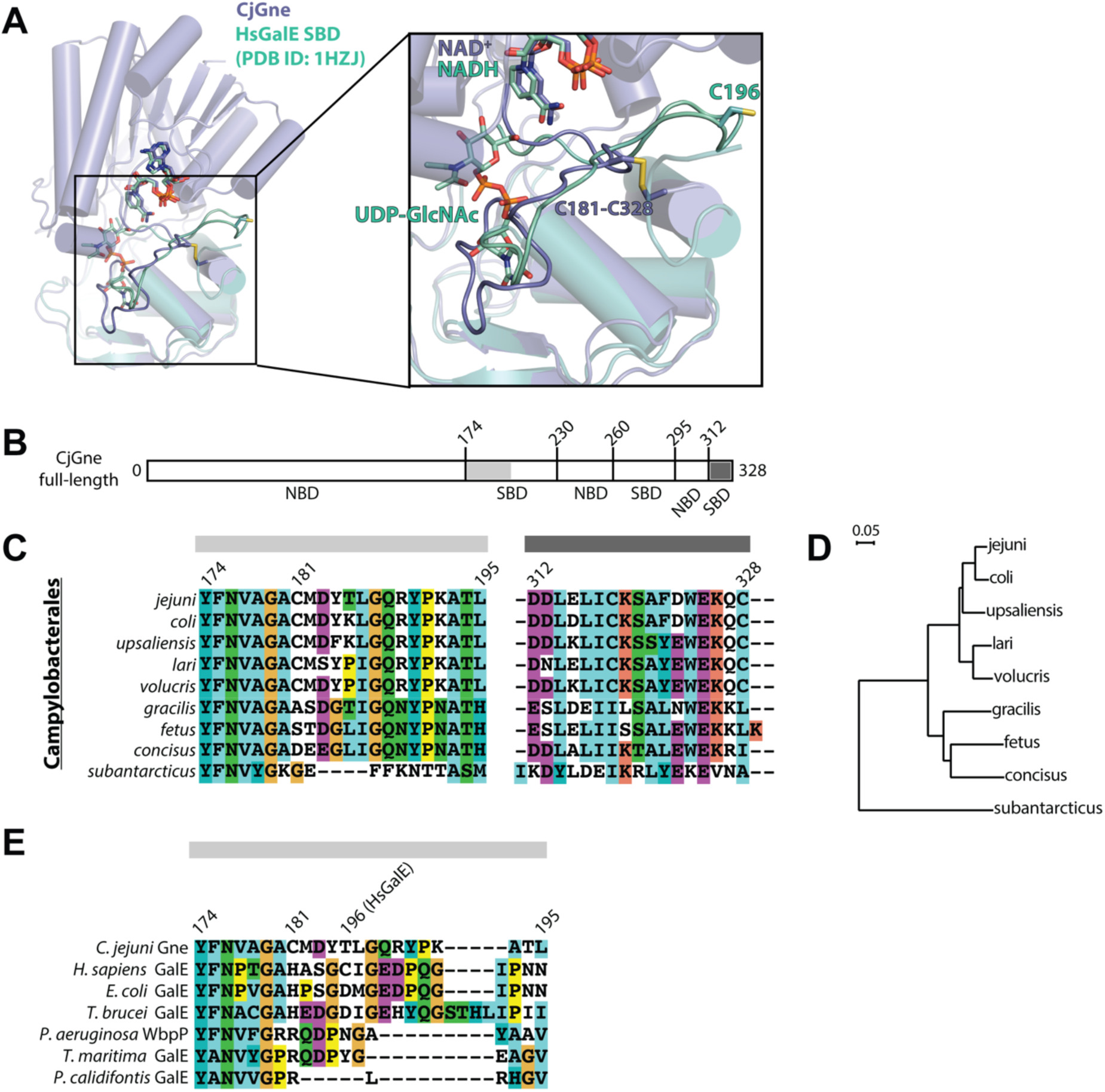
Unique features of *Cj*Gne. **A**. Structural alignment between *Cj*Gne/NAD^+^ (slate) and *Hs*GalE/NADH/UDP-GlcNAc (PDB ID: 1HZJ) (green cyan). The SBD is magnified in the box. NAD^+^, NADH, and UDP-GlcNAc are shown in the stick representation. C181 and C328 of *Cj*Gne along with the disulfide bond and C196 of *Hs*GalE are also in the stick representation. **B**. The full length of *Cj*Gne. **C**. Multiple sequence alignment of Campylobacterales using ClustalX 2.1 (60). Only two alignment regions (Y174-L195 and D312-C328; *Cj*Gne numbering) that are highlighted in the sequence scheme in figure 4B are shown. The background coloring is as follows: aromatic (cyan), hydrophobic (blue), polar (green), glycines (orange), negative charge (purple), positive charge (red), prolines (yellow), and unconserved (white). **D**. A phylogenetic tree of the Campylobacterales was drawn in NJplot (61). **E**. Multiple sequence alignment of *Cj*Gne and its homologs using ClustalX 2.1 (60). The alignment with only the region from Y174 to L195 (*Cj*Gne numbering) is shown here. The sequences that are used here are, from the top, Gne from *Campylobacter jejuni*, GalE from *Homo sapiens*, GalE from *Escherichia coli*, GalE from *Trypanosoma brucei*, WbpP from *Pseudomonas aeruginosa*, GalE from *Thermotoga maritima*, and GalE from *Pyrobaculum calidifontis*. The color scheme is the same as in Figure 4C. Please see Supplementary Figure 1 for the full-length multiple sequence alignment.

### The internal cysteine forming the disulfide bond is important for activity

We first examined the *Cj*Gne-catalyzed percent conversion of the four substrates (UDP-Glc, UDP-Gal, UDP-GlcNAc, and UDP-GalNAc) at equilibrium using capillary electrophoresis (CE). When the sample reaction started from either UDP-Glc or UDP-Gal, the ratio for the integral areas reached 23:75 (UDP-Glc:UDP-Gal) at equilibrium irrespective of the starting substrate (Figure 5A, 5B, and Table 2). Similarly, in the presence of either UDP-GlcNAc or UDP-GalNAc as a substrate, the reactions reached the ratio of 28:71 (UDP-GlcNAc:UDP-GalNAc) at equilibrium (Figure 5A, 5B, and Table 2). With this assay we confirmed the bi-functional activity of *Cj*Gne in interconverting similar amounts of non-acetylated and acetylated substrates.

**Table 2.** Average percent conversion values and their standard deviations of the substrates by *Cj*Gne and *Hs*GalE. The protein samples include the wild type (WT) and mutants. Ebselen, an inhibitor, was added only to *Cj*Gne. The normalized mean and standard deviation of percent conversion values of UDP-hexose substrates are shown (See **Experimental procedures** for the normalized standard deviation calculation).

| Enzymes | UDP-Glc |  | UDP-Gal |  | UDP-GlcNAc |  | UDP-GalNAc |  |
| --- | --- | --- | --- | --- | --- | --- | --- | --- |
|  | raw | normalized | raw | normalized | raw | normalized | raw | normalized |
| ***Cj*Gne (WT)** | **23.2 ± 0.2** | **100.0 ± 1.2** | **75.3 ± 1.0** | **100.0 ± 1.9** | **27.7 ± 0.2** | **100.0 ± 1.1** | **71.4 ± 0.7** | **100.0 ± 1.4** |
| C181A | 1.7 ± 0.5 | 7.5 ± 2.1 | 0.6 ± 0.3 | 0.7 ± 0.4 | 5.8 ± 0.2 | 20.8 ± 0.7 | 1.4 ± 1.2 | 2.0 ± 1.7 |
| C181S | 1.2 ± 0.1 | 5.3 ± 0.5 | 1.0 ± 0.4 | 1.4 ± 0.5 | 6.1 ± 0.5 | 22.1 ± 1.9 | 0.9 ± 0.8 | 1.3 ± 1.1 |
| C328S | 8.8 ± 3.2 | 37.9 ± 13.8 | 10.6 ± 3.9 | 14.0 ± 5.2 | 9.4 ± 0.4 | 34.0 ± 1.5 | 14.4 ± 3.3 | 20.1 ± 4.6 |
| ΔC328 | 23.6 ± 0.2 | 101.9 ± 1.2 | 76.0 ± 0.3 | 100.9 ± 1.4 | 27.5 ± 0.1 | 99.3 ± 0.9 | 72.1 ± 0.7 | 100.9 ± 1.4 |
| C181S/<br>ΔC328 | 1.1 ± 0.5 | 4.8 ± 2.3 | 0.6 ± 0.3 | 0.8 ± 0.4 | 5.9 ± 0.2 | 21.4 ± 0.9 | 0.7 ± 0.6 | 0.9 ± 0.8 |
| T122V<sup>c</sup> | 0 ± 0 | 0 ± 0 | 0 ± 0 | 0 ± 0 | 4.5 ± 2.9 | 16.3 ± 10.3 | 0 ± 0 | 0 ± 0 |
| N176L<sup>a</sup> | 0 ± 0 | 0 ± 0 | 0 ± 0 | 0 ± 0 | 0 ± 0 | 0 ± 0 | 0 ± 0 | 0 ± 0 |
| L294V<sup>c</sup> | 11.4 ± 5.2 | 49.4 ± 22.5 | 40.7 ± 11.9 | 54.0 ± 15.8 | 10.8 ± 2.0 | 39.0 ± 7.2 | 10.5 ± 6.9 | 14.7 ± 9.7 |
| L294M<sup>c</sup> | 2.7 ± 2.3 | 11.8 ± 9.7 | 12.1 ± 5.8 | 16.1 ± 7.7 | 12.1 ± 2.1 | 43.8 ± 7.5 | 12.5 ± 8.5 | 17.4 ± 11.9 |
| L294Y<sup>c</sup> | 2.4 ± 3.5 | 10.4 ± 15.3 | 10.6 ± 7.4 | 14.1 ± 9.9 | 5.8 ± 0.7 | 20.8 ± 2.5 | 0.4 ± 1.0 | 0.6 ± 1.4 |
| WT +<br>ebselen | 2.9 ± 1.8 | 12.6 ± 7.8 | 1.5 ± 2.1 | 2.0 ± 2.7 | 6.1 ± 0.1 | 22.0 ± 0.5 | 0 ± 0<sup>b</sup> | 0 ± 0 |
| ***Hs*GalE (WT)** | **25.2 ± 1.0** | **100.0 ± 5.7** | **72.6 ± 0.6** | **100.0 ± 1.2** | **23.3 ± 1.4** | **100.0 ± 8.5** | **56.3 ± 5.9** | **100.0 ± 14.9** |
| C196A | 25.1 ± 1.1 | 99.8 ± 5.9 | 62.8 ± 4.0 | 86.5 ± 5.6 | 18.9 ± 3.0 | 81.3 ± 13.9 | 47.9 ± 3.1 | 85.1 ± 10.5 |
| C196S | 25.7 ± 1.2 | 102.2 ± 6.4 | 68.4 ± 1.4 | 94.2 ± 2.1 | 19.7 ± 1.1 | 84.8 ± 6.9 | 52.0 ± 4.7 | 92.4 ± 12.8 |

The <sup>a</sup>, <sup>b</sup>, and <sup>c</sup> refers to a single, duplicate, and quintuplicate measurement, respectively. The rest of the values are from triplicates.

**Figure 5.**
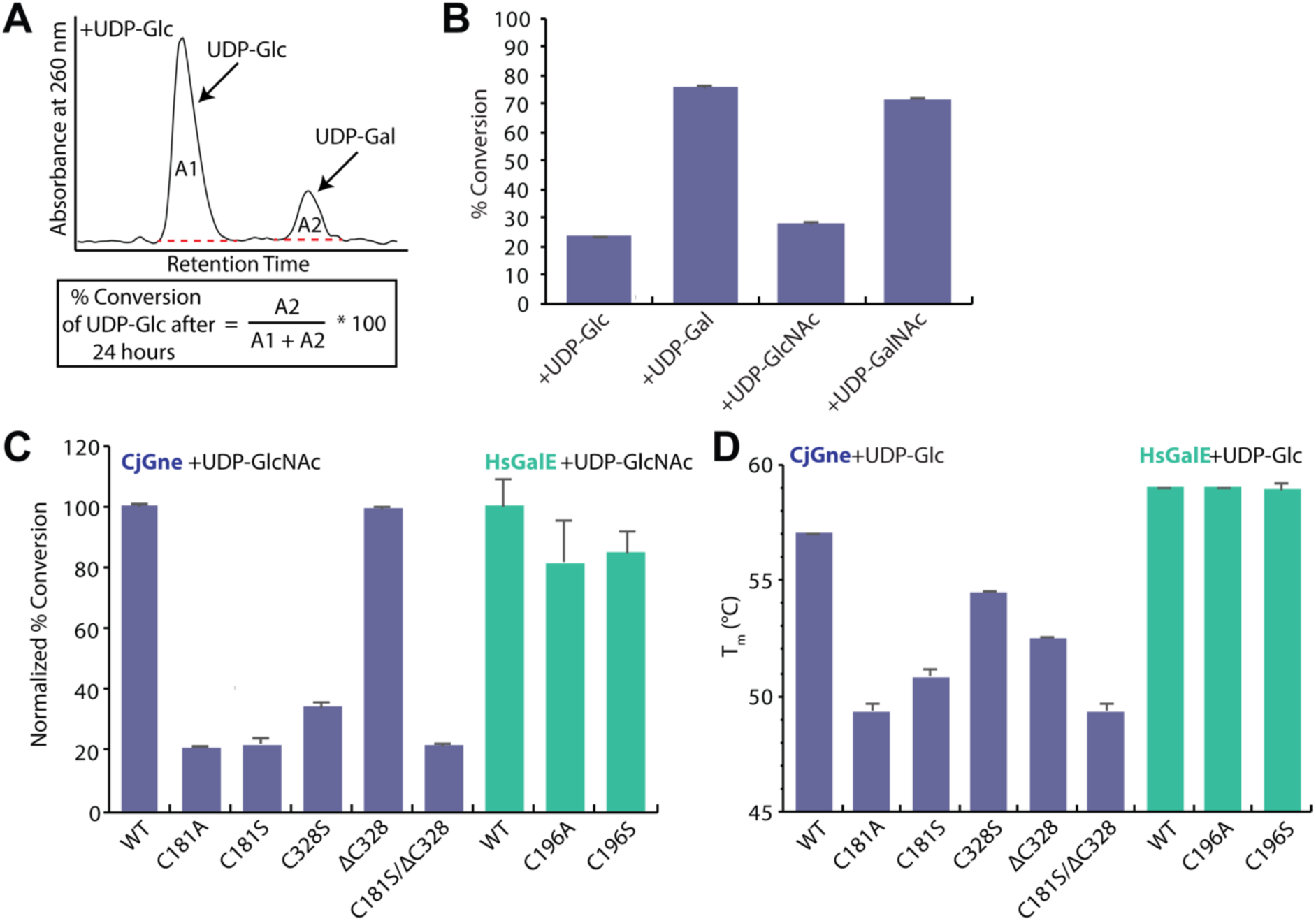
Epimerization assays of the wild type and cysteine mutants of *Cj*Gne and *Hs*GalE. **A**. An example of the % conversion of UDP-Glc to UDP-Gal by *Cj*Gne measured in capillary electrophoresis. The equation used to calculate % conversion is provided in the box. **B**. The results of % conversion of each of four substrates by *Cj*Gne. Each reaction was repeated for three times and the error bars indicate the standard deviations. **C**. Normalized % conversion of UDP-GlcNAc by the wild type and cysteine mutants of *Cj*Gne (slate) and *Hs*GalE (green cyan). Each reaction was repeated for three times and the error bars are the standard deviations (See **Experimental procedures** for the normalized standard deviation calculation). **D**. The melting temperature values (T_m_) of the wild type and cysteine mutants of *Cj*Gne (slate) and *Hs*GalE (green cyan) in the presence of UDP-Glc. Each measurement was repeated for three times and the error bars indicate the standard deviations.

The unexpected disulfide bond brings into question whether there may be biological importance to these residues. To examine the roles of C181 and C328, we expressed and purified various cysteine mutants (C181A, C181S, C328S, ΔC328, and C181S/ΔC328) and analyzed their activities at equilibrium (Figure 5C and Table 2). All of the C181 mutants (C181A, C181S, and C181S/ΔC328) resulted in significant loss of activity, less than 20% of wild type conversion, irrespective of the starting UDP-hexose. For the C-terminal cysteine, the C328S mutant lost more than 50% of the wild-type conversion. Yet surprisingly, the deletion of C-terminal C328 retained the full activity of the wild type regardless of the substrate added.

### The similar *Hs*GalE cysteine is not required for activity

In the multiple sequence alignment of *Cj*Gne and its homologs (Figure 4E), there is no epimerase that has a cysteine residue at the corresponding position to C181 of *Cj*Gne. Instead, the human epimerase among the homologs is the only example that has a cysteine residue (C196) within the corresponding loop region (Y185-N207) to the previously mentioned loop (Y174-L195) in *Cj*Gne. We expressed and purified the C196A and C196S mutants of *Hs*GalE and investigated their epimerization activity at equilibrium in the reaction with each of the sugar substrates. Again, we confirmed the bifunctional nature of the enzyme. In this case, both mutants had no significant loss in activity relative to the wild type (Figure 5C and Table 2).

### The *Cj*Gne internal cysteine is important in thermal stability of the enzyme

Thermal denaturation of the wild-type *Cj*Gne and its cysteine mutants was monitored through melting curves and calculated melting temperature (T_m_) (Figure 5D and Table 3). The T_m_ values across the four UDP-hexose substrates within an enzyme stayed the same or had a difference of less than 1°C. All the cysteine mutants of *Cj*Gne had T_m_ values that are lower than the wild type by 1°C or more. The single and double mutants containing C181A decreased the T_m_ values the most by 8°C. Protein unfolding was also monitored for the *Hs*GalE C196 mutants (Figure 5D and Table 3). Unlike those of the *Cj*Gne mutants, the T_m_ values did not change.

**Table 3.** Average melting temperatures (°C) and their standard deviations of the wild type (WT) and mutants of *Cj*Gne and *Hs*GalE*.

| Enzymes | Melting Temperature (°C) |  |  |  |
| --- | --- | --- | --- | --- |
|  | UDP-Glc | UDP-Gal | UDP-GlcNAc | UDP-GalNAc |
| *CjGne* (WT) | 57.0 | 57.2 ± 0.3 | 58.5 | 58.0 |
| C181A | 49.3 ± 0.3 | 49.5 | 51.3 ± 0.3 | 50.5 |
| C181S | 50.8 ± 0.3 | 50.7 ± 0.3 | 53.0 ± 0.5 | 51.5 |
| C328S | 54.5 | 54.2 ± 0.3 | 55.5 | 55.0 |
| ΔC328 | 52.5 | 52.0 | 53.8 ± 0.6 | 52.5 |
| C181S/ΔC328 | 49.3 ± 0.3 | 49.5 | 50.2 ± 0.3 | 50.2 ± 0.3 |
| *HsGalE* (WT) | 59.0 | 59.0<sup>a</sup> | 59.0 | 59.0 |
| C196A | 59.0 | 59.0 | 59.0 | 59.0 |
| C196S | 59.0 ± 0.2 | 59.0 | 59.0 | n.d. |

The <sup>a</sup> refers to a single measurement. The rest of the values are from at least duplicates.
\* Standard deviations of zero are not shown.

### Critical residues of *Cj*Gne for substrate binding and specificity

We performed a structural alignment between *Cj*Gne and *Hs*GalE bound with UDP-GlcNAc (PDB ID: 1HZJ) (Figure 6A). From the alignment, we identified five residues of *Cj*Gne that can make polar contacts with the UDP-GlcNAc (T122, N176, P185, K192, and T194). Among them, both T122, either a serine or threonine, and N176 is conserved among the homologous sequences (Figure 4E and Supplementary Figure 1). In order to examine their functional roles within the predicted active site, we introduced mutations that retain the size, but lose polarity of the side chain: T122V and N176L (Figure 6A).

**Figure 6.**
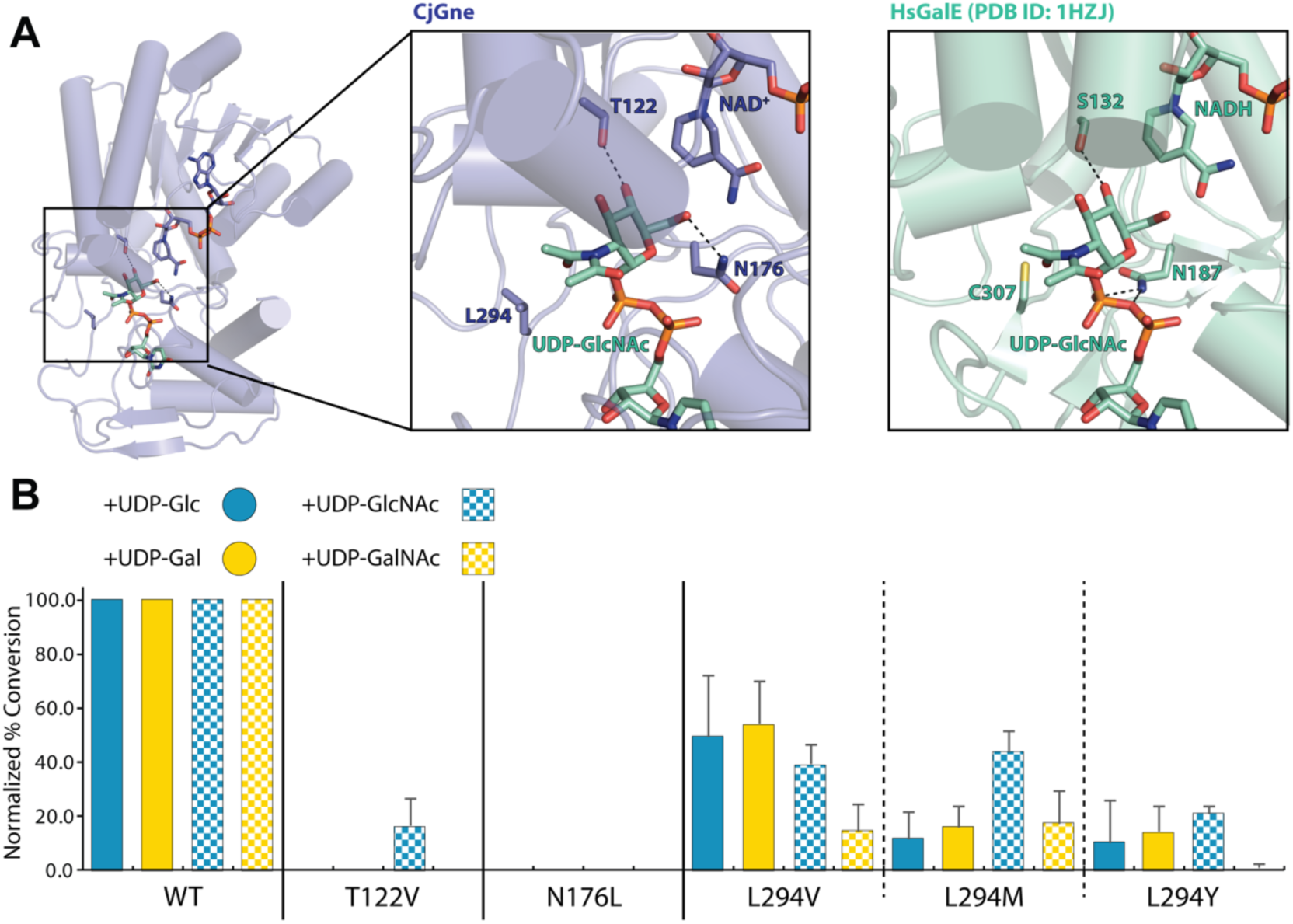
Predicted active site of the *Cj*Gne/NAD^+^ complex. **A**. On the left, structural alignment between *Cj*Gne/NAD^+^ and UDP-GlcNAc from the *Hs*GalE/NADH/UDP-GlcNAc (PDB ID: 1HZJ). In the center, the putative active site of *Cj*Gne is magnified with T122, N176, and L294 labeled. On the right, the catalytic site of *Hs*GalE is magnified to the same extent as that with *Cj*Gne with S132, N187, and C307 labeled. Hydrogen bonds are indicated as black dashes. **B**. Normalized % conversion of UDP-Glc, UDP-Gal, UDP-GlcNAc, and UDP-GalNAc by the wild type and catalytic mutants of *Cj*Gne. Each reaction was repeated for five times except for N176L only for once. The error bars are the standard deviations (See **Experimental procedures** for the normalized standard deviation calculation).

Based on the classification scheme for UDP-hexose 4-epimerases (33), the T122 and N176 residues are two of the six key residues in *Cj*Gne, determining substrate specificity. Another key residue, L294, was also subject to mutation varying the size of hydrophobic side chains (valine, methionine, and tyrosine) to investigate its role in substrate specificity (Figure 6A). The corresponding residue of L294 in *Ec*GalE (Y299) was the first example that showed a single-residue mutation at this position to cysteine altered substrate specificity (39).

The equilibrium assay was then performed with each of the mutants. For T122V and N176L, there was nearly a complete loss of conversion for all of the four UDP-sugar substrates (Figure 6B). This supports that T122 and N176 have catalytic roles that are essential for the epimerization activity of *Cj*Gne. T122 of *Cj*Gne is structurally aligned to S124 (*Ec*GalE), which is suggested to mediate catalysis by being hydrogen-bonded to Y149 (*Ec*GalE), an active site base, and to the 4’-hydroxyl group of the glucosyl ring of UDP-Glc (40). Also, in *Hs*GalE, the corresponding residue, S132 interacts with the 4’-OH group of hexose. In *Cj*Gne, T122 and Y146 (Y149 in *Ec*GalE) are not in hydrogen-bonding distance.

N179 of *Ec*GalE, N176 in *Cj*Gne, has been suggested as one of the amino acid residues that contact the hexose portion of UDP-Glc/Gal within a hydrogen-bonding distance and thereby important for binding of the substrate in the *E. coli* (27). *Cj*Gne N176 is in hydrogen bonding distance of the modeled UDP-GlcNAc (Figure 6A). Alternatively, the amino group of N187 (*Hs*GalE) interacts directly with the two phosphoryl oxygen atoms of the β-phosphate group of the nucleotide. With substrate bound N176 may re-orient in *Cj*Gne to make this same interaction.

The epimerization assay was also performed for the L294V, L294M, and L294Y mutants to investigate if the size of the L294 side chain alters substrate specificity of *Cj*Gne (Figure 6B). None of the mutants retained activities at a comparable level to those of the wild type with all four substrates. However, among the mutants L294V had the least disruption for interconversion of non-acetylated substrates, whereas L294M had the highest activity for acetylated substrates.

### Ebselen inhibiting *Hs*GalE is a potential inhibitor of *Cj*Gne

Few studies have reported inhibitors of GalE, and none of them are promising for further drug development. One report identified ebselen, an organoselenium compound, inhibition of *Hs*GalE with nanomolar IC_50_ (0.014 μM) (41). Since *Cj*Gne possesses a bifunctional activity just like *Hs*GalE, we predicted that the inhibitors that are potent to *Hs*GalE would also exert inhibitory effect on *Cj*Gne. An inhibition assay of *Cj*Gne using capillary electrophoresis was performed in the presence of ebselen (Figure 7). Indeed, the inhibition was almost complete when ebselen was incubated with the enzyme with either UDP-Gal or UDP-GalNAc, while slightly less inhibition for UDP-Glc or UDP-GlcNAc as initial substrates (13% and 22% of the wild-type activity was retained, respectively).

**Figure 7.**
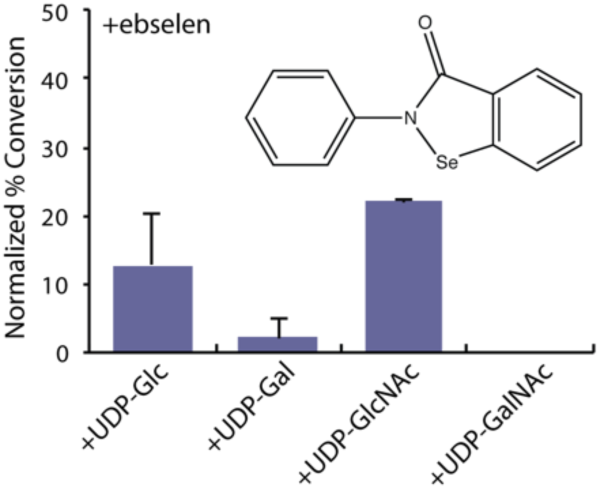
Inhibition of *Cj*Gne by ebselen. Chemical structure of ebselen is shown in the top right corner. Each inhibition reaction was repeated for three times except for the one with UDP-GalNAc only for twice. The error bars are the standard deviations (See **Experimental procedures** for the normalized standard deviation calculations).

## Discussion

In this study, we provide the first structure and biochemically characterize *Cj*Gne, a bifunctional UDP-Glc/GlcNAc 4-epimerase, the sole enzyme producing the GalNAc residue for the major surface polysaccharides in *C. jejuni*. Gne is functionally separated from other forementioned UDP-Glc/GlcNAc 4-epimerases that either work in the Leloir pathway for galactose metabolism or are part of the LPS biosynthesis pathway.

A surprising result is the presence of a structural disulfide bond in *Cj*Gne. Expressed heterologously in the *E. coli* system, in the cytosol one expects these cysteines to be reduced (42). Other exceptions have been reported where structural disulfides in cytosolic proteins of thermophilic archaeal species protect them from denaturation at high temperature (43). *C. jejuni* is a moderate thermophilic species thriving at 37-42°C (44) with a T_m_ for *Cj*Gne at 57°C (Figure 5D and Table 3). The presence of a structural disulfide bond might help *Cj*Gne tolerate growing at higher temperature.

The cysteine at position 181 is not conserved (Figure 4E), yet replacing C181 with either alanine or serine led to a loss of activity. In the structure, a water mediated network connects the phosphate groups of the substrate to the backbone of the C181 residue. These are unlikely to be disrupted by the simple mutations. When the other half of the disulfide pair, C328, was replaced with a serine, less than 50% wild-type conversion was retained. Surprisingly, deletion of the C-terminal cysteine (C328) was fully active. In total, we can conclude that the disulfide bond is not required for the catalysis. However, the cysteines are important for full activity. Cysteines often make hydrophobic contacts and the effects from the loss of the spatial hydrophobic environment may provide rationale for the reduction in activity upon mutation to serine. Supporting this, all the cysteine mutants of *Cj*Gne reduced the overall stability of the protein (Figure 5D and Table 3). This suggests general structural roles that could affect catalysis.

A major feature of our *Cj*Gne structure is the shift of the loop region (Y174-L195) toward the active site in *Cj*Gne. The presence of the disulfide suggests that this may fix the conformation of the loop. The sequence of this region is conserved across other *Campylobacter* species and the electron density shows the conformation is well ordered. In this conformation, the protein would be expected to be inhibited. We can speculate on a biological role for this cysteine mediated shift. It had previously been shown that the mucosal defense of the release of reactive oxygen species led to inactivation of *Cj*Gne lowering virulence and altering surface polysaccharides (45). Here it was shown also that an important population of *Cj*Gne localized to the outer membrane. *C. jejuni* regulates cell envelope genes, including Gne, during infection (46). These various points support that Gne function might be modulated as a response to various cellular environments, perhaps to facilitate infection under stress conditions.

While the cysteines are important for functional protein in *C. jejuni*, a related cysteine in the functional homolog *Hs*GalE was not required for full activity. In the structure of the human there is no shift of the corresponding *Cj*Gne loop (Y174-L195). This lends credence to the importance of these residues in the *C. jejuni* enzyme.

The mechanism of this class of epimerases has been elucidated over time; however, there remain some details to be resolved. We chose candidate residues that are predicted to be critical in substrate binding or specificity from the structural and sequence alignments with the human epimerase in complex with NADH and UDP-GlcNAc (PDB entry: 1HZJ). Both the T122V and N176L mutants lost activity irrespective of the substrate tested. This result is indicative of catalytic importance of the residues. The highly conserved serine residues of the other homologs aligned with T122 of *Cj*Gne have been shown to play a role in mediating the electron transfer during the catalytic mechanism. The asparagine residue corresponding to N176 in *Cj*Gne is also almost perfectly conserved across species. The N195 residue in the *Ps*WbgU (N176; *Cj*Gne), for example, forms two hydrogen bonds via the carbonyl oxygen and the amine group with the NH-group and the oxybridge of the diphospho moiety of UDP-GlcNAc, respectively. Based on the high conservation and previous functional characterization in other homologs, T122 and N176 of *Cj*Gne are likely to require polar side chains to be either directly or indirectly involved in catalysis and to interact with the substrate bound, respectively.

L294 of *Cj*Gne was chosen for mutagenesis studies because it had previously been shown for homologs that mutations at this position altered substrate specificity. Previous studies demonstrated that Y299C of *Ec*GalE (47) allowed conversion of acetylated substrates, while C307Y of *Hs*GalE (48) and S306Y of *Ec*GalE_O86:B7_(49) lost the conversion activity of acetylated substrates. In contrast, mutation of the corresponding residue from *Pa*WbpP (S317Y) resulted in complete loss of activity, so no further insights regarding side chain size of this residue for substrate specificity was available (33). We mutated the residue in the same position, L294, of *Cj*Gne into valine, methionine, and tyrosine to see if various sizes of hydrophobic side chains can affect the activities of any UDP-hexose substrate. We predicted that L294M would retain bifunctionality, whereas L294V and L294Y would preferentially convert the acetylated and non-acetylated sugar substrates, respectively. None of the three mutants of L294 retained the full activities of the wild type, but showed similar levels of reduction between UDP-Glc and UDP-Gal and generally more reduction with UDP-GalNAc. Approximately 50% reduction in L294V activity with non-acetylated substrates can be explained by the smaller hydrophobic side chain of valine that increased the active site volume, rendering the binding to non-acetylated and/or acetylated substrates less specific. Also, the hydrophobic interaction between leucine and the methyl group of the N-acetylated moiety on C2 on UDP-GlcNAc from the human enzyme seems to be disrupted in *Cj*Gne by the replacement with valine, thereby resulting in a larger reduction in conversion activity of the acetylated substrates. Methionine is similar in size to leucine and the L294M mutation had similar levels of activity for both non-acetylated and acetylated substrates except for UDP-GlcNAc. In all cases, the epimerization is much less efficient than that catalyzed by the wild type. For the L294Y mutant, the bulky side chain of tyrosine likely reduced the active site volume, thus leading to the highest loss of epimerization with UDP-GalNAc. Taken together, we showed the variants of single active site residue (L294) could alter substrate specificity, although the previously predicted pattern did not fit here.

Lastly, the inhibition of *Cj*Gne by ebselen provides evidence for this as a small molecule target, but ebselen itself is not a likely route as it has been reported to target myriads of biological pathways (50). It will be important to find inhibitors that are specific to *Cj*Gne. The search for such inhibitors is ongoing.

In conclusion, based on a high-resolution *Cj*Gne/NAD^+^ crystal structure and biochemical data from mutants, we proposed some critical residues of *Cj*Gne that are catalytically or structurally important. These findings along with its observed susceptibility to a *Hs*GalE inhibitor, suggest a route for antibiotic development.

### Experimental procedures Materials and chemicals

Genomic DNA of *C. jejuni* NCTC11168 (#700819D-5) was purchased from ATCC (Manassas, VA). Two constructs of *Hs*GalE with either N-terminal (pET28a vector) or C-terminal (pET31B vector) hexahistidine (His_6_) tags were kindly provided by the Holden Group at the University of Wisconsin, Madison. The former was used for capillary electrophoresis and the latter was used for stability assays, each described below. *E. coli* NiCo21(DE3) competent cells were obtained from New England Biolabs Inc. (Ipswich, MA). UDP-Glucose, UDP-Galactose, UDP-GlcNAc, UDP-GalNAc, nicotinamide adenine dinucleotide (NAD^+^), dithiothreitol (DTT), tris (2-carboxyethyl) phosphine (TCEP), and bovine serum albumin (BSA) were from Sigma-Aldrich (St. Louis, MO). Ebselen (2-phenyl-1,2-benzisoselenazol-3(2H)-one) was from Cayman Chemical (Ann Arbor, MI).

### Cloning of gne

The plasmid encoding *gne* (*cj1131c*) of *C. jejuni* was constructed by one-step enzymatic DNA assembly through Gibson cloning (51). Briefly, the *gne* gene and the plasmid for insertion were amplified from genomic DNA of *C. jejuni* NCTC11168 and a pET33b-derived vector, respectively, using primers with ca. 40 bp of homology to each other. 5 μL of DNA sample (0.7 μL of the amplified *gne* gene (10 ng/μL) plus 4.3 μL of the amplified vector (10 ng/μL)) was added to 15 μL of a master mix solution including T5 exonuclease, Phusion DNA polymerase and Taq DNA ligase, then incubated at 50°C for 60 min. To avoid self-colonies from the template-vector, the PCR product amplified from the vector was treated with *DpnI*. The final construct contained a N-terminal His_6_-tag followed by a thrombin cleavage site (N-**M**GG-S**HHHHHH**G**LVPRGS**-*gne*-C). All DNA constructs were confirmed by sequencing.

### Generation of point mutations

All mutations, including T122V, N176L, L294V, L294M, L294Y, C181A, C181S, C328S, ΔC328, C181S-ΔC328, C196A (*Hs*GalE), and C196S (*Hs*GalE) were prepared in a mixture solution containing Phusion^®^ High-Fidelity PCR Master Mix with HF Buffer from New England Biolabs Inc. (Ipswich, MA), the DNA template of the QK45AA mutant (based on the surface entropy reduction prediction (52)), and primers with or without 5% DMSO (53). The recommended PCR protocol for using the Phusion^®^ HF PCR MM from NEB Inc. was used.

### Expression and purification of *Cj*Gne

Constructs were transformed and expressed in *E. coli* NiCo21(DE3) strain that are originally derived from BL21(DE3). Cells were grown in cultures of 2xYT with kanamycin (35 μg/mL) to an optical density of OD_600_ 0.5-0.7, then induced with 0.4 mM isopropyl β-D-thiogalactoside (IPTG, Anatrace, Maumee, OH) at 37°C for 6 h. Cells were harvested by centrifugation (4,000 rpm, 20 min, 4°C), resuspended in lysis buffer (20 mM Tris-HCl, pH 7.5, 100 mM NaCl, 1 mM DTT), and lysed by three passes through a microfluidizer (18 kpsi), and centrifuged at 12,000 rpm for 20 min at 4°C. Proteins were purified from the supernatant by nickel affinity chromatography followed by size exclusion chromatography. Briefly, the cell lysate was modified to 10 mM imidazole then passed through a 1 mL pre-equilibrated Ni-NTA agarose resin (Qiagen, Germantown, MD), washed with 30x column volumes of 20 mM imidazole-containing buffer, then eluted with 20 mL of 250 mM imidazole-containing buffer. The elution was concentrated using a 10-kD cut-off Amicon^®^ Ultra-4 Centrifugal Filter (Millipore) and further purified by either a Super-dex-200 16/60 or Superdex-200 10/300 gel-chromatography column (GE Healthcare, Little Chalfont, UK). The fractionated protein was concentrated again using the Amicon^®^ Centrifugal Filter, flash frozen, and stored at -80°C. In all cases, point mutants behaved similar to wild-type during purification.

### Size exclusion chromatography and multi-angle laser light scattering (SEC-MALLS)

Purified protein was analyzed by SEC-MALLS (Wyatt Technology, Santa Barbara, CA). Briefly, a Shodex KW-804 column (Showa Denko America, Inc., New York, NY) was equilibrated in running buffer (20 mM HEPES, pH 7.5, 100 mM NaCl, 10% glycerol, 10 mM βME). 100 μg of either BSA as a standard or *Cj*Gne were injected and run at 0.5 mL/min for 30 min. Data analysis was performed using Astro 5.3.4 software. For the sample run, peaks from LS, dRI, and UV (280 nm) detectors were aligned by defining baseline and applying band broadening values from the BSA run.

### Crystallization of *Cj*Gne and x-ray diffraction

For crystallization, the purified QK45AA optimized variant of *Cj*Gne was pre-incubated with 5 mM UDP-GlcNAc at 4°C overnight. Crystallization screening was performed by sitting drop vapor-diffusion with commercially available screens (Hampton Research, Qiagen, Emerald BioSystems) and then incubated at room temperature. Initial conditions were refined by additive screening using the Additive Screen^™^ (Hampton Research). The final drop consisted of 0.2 μL of mother liquor (1.3 M sodium acetate trihydrate (pH 7.0) with 50 mM sodium malonate) and 0.2 μL of protein (16-20 mg/mL). Crystals grew to full-size after several days. For cryo-protection, crystals were transferred to a drop containing 70% reservoir solution and 30% glycerol for 5 sec then flash frozen in liquid nitrogen. Diffraction data were collected from a single crystal at beamline 12-2 at the Stanford Synchrotron Radiation Lightsource (SSRL). Despite numerous attempts, we were never able to obtain crystals of our point-mutants with or without the QK45AA mutation.

### Structural determination and refinement

Images were collected on a Dectris Pilatus 6M pixel detector. Diffraction data were integrated with XDS (54) and scaled with SCALA in CCP4 (55). Crystals were in the space group P4_1_2_1_2 with unit cell dimensions *a=b=87*.*7 c=261*.*66* and a complete dataset was collected to 2.0 Å. The asymmetric unit contained two copies of *Cj*Gne (residues 2-328), two NAD^+^, two acetate, seven glycerol, and 167 water molecules. Phases were obtained by molecular replacement using the structure of UDP-Glc 4-epimerase from *Bacillus Anthracis* as a search model (PDB entry: 2C20; 42% identity) in Phaser as implemented in Phenix (56, 57). Manual model building was performed using Coot (58). *Cj*Gne was refined in Phenix with final R-factor of 19.5% (R_free_ = 22.5%). Statistics for data collection and structure determination are found in Table 1.

### Epimerization assay using capillary electrophoresis

Enzyme reactions were performed in 100 μL of reaction mixture containing 50 mM Tris-HCl (pH 8.0), 1 mM of UDP-sugar, 1 mM of NAD^+^, and 50 ng of *Cj*Gne at 37°C. The reaction was stopped after 24 hours by boiling for 5 min and then centrifuged (12,000 rpm, 20 min) to remove protein aggregates. In the case of the inhibition assay with ebselen, enzyme reactions were prepared in 20 μL containing 50 mM Tris-HCl (pH 8.0), a 1 mM of UDP-sugar, 1 mM of NAD^+^, 50 ng of *Cj*Gne, and 100 μM ebselen at 37°C for 24 hours. The reactions were then stopped by boiling for 5 min and centrifuged. The samples were monitored using a HP 3DCE capillary electrophoresis instrument equipped with a UV-VIS diode-array detector (DAD) (Agilent Technologies, Santa Clara, CA, USA). The 50 cm-long capillary was packed with fused silica and the running buffer was 20 mM sodium tetraborate decahydrate, pH 9 (J.T. Baker®, Avantor Performance Materials, Center Valley, PA). The capillary was preconditioned for each run by washing with the running buffer for 2 min. Each sample was injected by pressure of 50 mbar for 10 sec and the separation was performed at 30 kV and detected at 260 nm (330 nm as background). The average retention times for UDP-Glc, UDP-Gal, UDP-GlcNAc, and UDP-GalNAc were 8.7 min, 8.9 min, 8.3 min, and 8.5 min, respectively. The peak area was estimated and integrated using the software 3D-CE Chemstation Rev. A.09.03.

Standard deviations (*σ*_*z*_) for the normalized conversion values in Table 2 and Figures 5-7 were calculated based on the equation below (*x*_0_: mean raw conversion value for mutant; *y*_0_: mean raw conversion value for wild type; *σ*_*x*_: standard deviation of raw conversion value for mutant; *σ*_*y*_: standard deviation of raw conversion value for wild type).

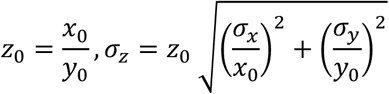

### A fluorescence-based thermal shift assay

The thermal shift assay was based on Niesen *et al* (59). A real-time PCR device (CFX96 from Bio-Rad, Hercules, CA) was used to monitor protein unfolding through fluorescence by SYPRO Orange (Sigma-Aldrich, St. Louis, MO) at 2x concentration (1:2,500 dilution of 5,000x stock). Fluorescence (λ_ex_ 470 nm/ λ_em_ 570 nm) was measured using the filter sets found on the real-time PCR instrument. Protein samples (2 μM) in 20 mM Tris buffer (pH 7.5) containing 100 mM NaCl, 1 mM NAD^+^, 1mM DTT, 1mM of a UDP-sugar substrate in a reaction volume of 50 μL were mixed in 96-well PCR plates (Bio-Rad, Hercules, CA). The plates were briefly spun down at 1,000 rpm and placed in the device. Melting curves were measured starting with a 15-minute pre-chilling at 15°C then increased in 0.5°C steps to 95°C with 30 second incubation. The fluorescence intensity at the end of each step is plotted as a function of temperature. The resulting sigmoidal curve was best fit to a two-state transition.

The inflection point of the melting curve, melting temperature (T_m_), was calculated using the internal PCR software.

## Data Availability

The structure presented in this paper has been deposited in the Protein Data Bank (PDB) with the following code: 7K3P. All remaining data are contained within the article.

## Acknowledgments

We thank Hazel Holden (Wisconsin) for expression plasmids for *Hs*GalE. We are grateful to Nathan Dalleska (Caltech) for assistance with the capillary electrophoresis at the Environmental Analysis Center and Jens Kaiser and Pavle Nikolovski (Caltech) for crystallography help through the Molecular Observatory.

## Funding and additional information

This work was supported by grants from the National Institutes of Health (NIH) National Institute of General Medicine (NIGMS) awards GM105385 and GM114611 (to WMC). This research was undertaken in part using the 12-2 beamline at the Stanford Synchrotron Radiation Lightsource (SSRL). Operations at SSRL are supported by the US Department of Energy and the National Institutes of Health (NIH).

---

**Supplementary Figure 1.**
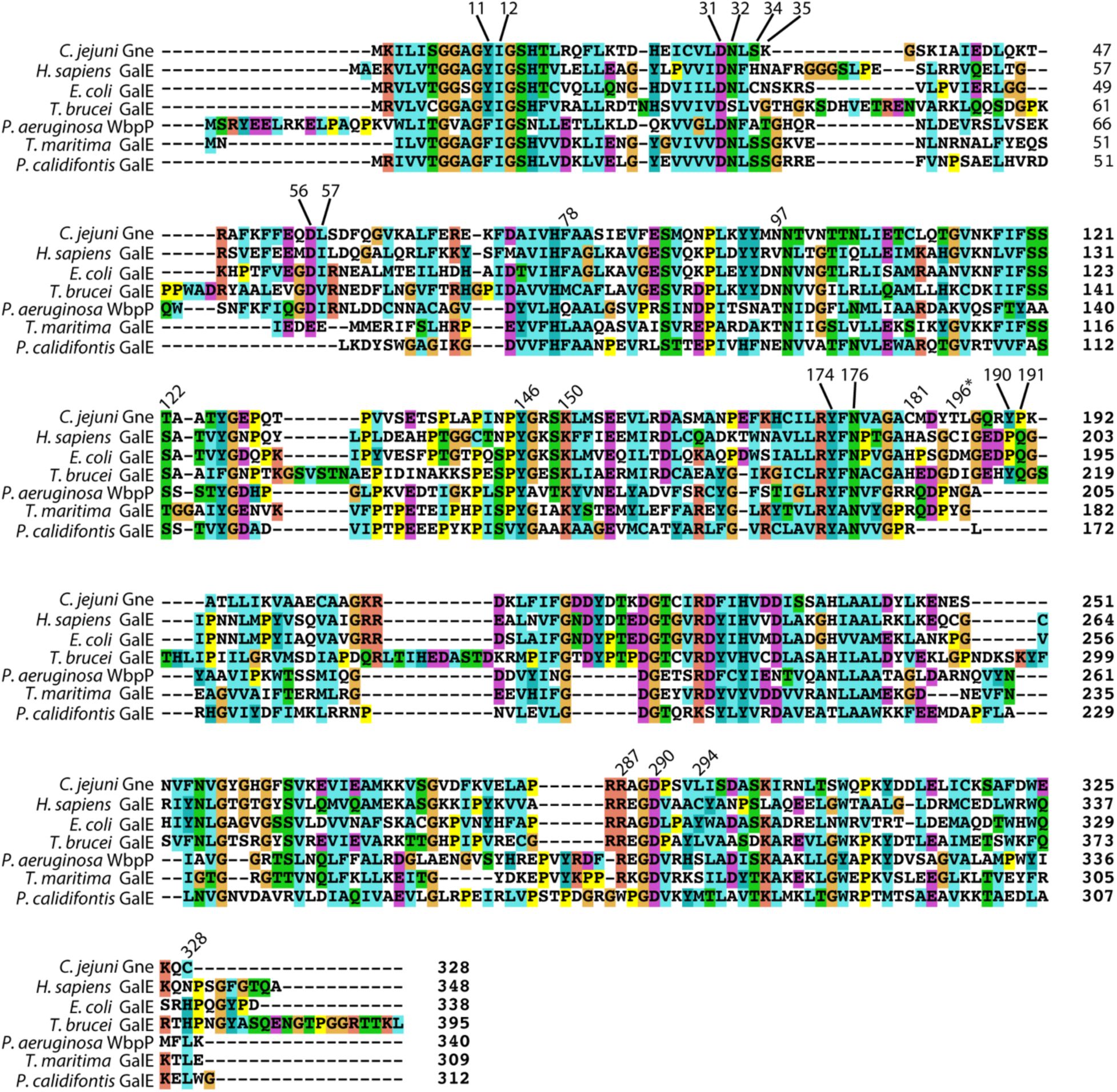
Multiple sequence alignment of the full-length *Cj*Gne and its homologs using ClustalX 2.1 (60). The sequences aligned are the same as in Figure 4E. The background coloring is the same as in Figure 4C and 4E. The numbers on the right indicate the last residue number of each segment of protein sequences. The numbers on the top of the segments, except for 196*, indicate the *Cj*Gne residues that are mentioned in this work. 196* indicates the cysteine residue in *Hs*GalE.

